# Dissecting the Causal Mechanism of X-Linked Dystonia-Parkinsonism by Integrating Genome and Transcriptome Assembly

**DOI:** 10.1101/149872

**Authors:** Tatsiana Aneichyk, William.T. Hendriks, Rachita Yadav, David Shin, Dadi Gao, Christine A. Vaine, Ryan L. Collins, Alexei Stortchevoi, Benjamin Currall, Harrison Brand, Carrie Hanscom, Caroline Antolik, Marisela Dy, Ashok Ragavendran, Patrick Acuña, Criscely Go, Yechiam Sapir, Brian J. Wainger, Daniel Henderson, Jyotsna Dhakal, Naoto Ito, Neil Weisenfeld, David Jaffe, Nutan Sharma, Xandra O. Breakefield, Laurie J. Ozelius, D. Cristopher Bragg, Michael E. Talkowski

**Affiliations:** Center for Genomic Medicine, Massachusetts General Hospital, Boston, MA, 02114, USA; Department of Neurology, Massachusetts General Hospital and Harvard Medical School, Boston, MA, 02114, USA; Program in Medical and Population Genetics and Stanley Center for Psychiatric Research, Broad Institute, Cambridge, MA, 02142, USA; The Collaborative Center for X-linked Dystonia Parkinsonism, Massachusetts General Hospital, Charlestown, MA, 02129, USA; Harvard Brain Science Initiative, Harvard Medical School, Cambridge, MA, 02138, USA; Program in Bioinformatics and Integrative Genomics, Division of Medical Sciences, Harvard Medical School, Boston, MA, 02115, USA; Jose Reyes Memorial Medical Center, Manila, 1003, Philippines; Genome Sequencing and Analysis Program, Broad Institute, Cambridge, MA, 02142, USA; Department of Radiology, Massachusetts General Hospital, Boston, MA, 02114, USA; Departments of Psychiatry and Pathology, Massachusetts General Hospital, Boston, MA, 02114, USA; Psychiatric and Neurodevelopmental Genetics Unit, Department of Psychiatry, Massachusetts General Hospital, Boston, MA, 02114, USA

**Keywords:** XDP, DYT3, SVA, retrotransposon, dystonia, Parkinson’s disease, *TAF1*, genome assembly, transcriptome assembly, intron retention

## Abstract

X-linked Dystonia-Parkinsonism (XDP) is a Mendelian neurodegenerative disease endemic to the Philippines. We integrated genome and transcriptome assembly with induced pluripotent stem cell-based modeling to identify the XDP causal locus and potential pathogenic mechanism. Genome sequencing identified novel variation that was shared by all probands and three recombination events that narrowed the causal locus to a genomic segment including *TAF1*. Transcriptome assembly in neural derivative cells discovered novel *TAF1* transcripts, including a truncated transcript exclusively observed in probands that involved aberrant splicing and intron retention (IR) associated with a SINE-VNTR-Alu (SVA)-type retrotransposon insertion. This IR correlated with decreased expression of the predominant *TAF1* transcript and altered expression of neurodevelopmental genes; both the IR and aberrant *TAF1* expression patterns were rescued by CRISPR/Cas9 excision of the SVA. These data suggest a unique genomic cause of XDP and may provide a roadmap for integrative genomic studies in other unsolved Mendelian disorders.

**Highlights:**

- Genome assembly narrows the XDP causal locus to a segment including *TAF1*
- XDP-specific SVA insertion induces intron retention and down-regulation of *TAF1*
- CRISPR/Cas9 excision of SVA rescues aberrant splicing and cTAF1 expression in XDP
- Gene networks perturbed in proband cells associate to synapse and neurodevelopment

## Introduction

In recent years, remarkable progress has been made in Mendelian gene discovery and characterization of the diverse phenotypic effects associated with disruption of normal gene function. Yet at present more than half of individuals with suspected genetic disorders do not receive a diagnosis, with recent estimates suggesting that ~20% of Mendelian disorders have been mapped to a causal locus but the pathogenic mechanism remains unknown (Chong et al., 2015; Lee et al., 2014; Yang et al., 2014). Critical limitations that continue to impede gene discovery in Mendelian disorders include the immature functional annotation of most coding and noncoding variation, as well as an inability to routinely survey all classes of structural rearrangements. Another barrier for genome and transcriptome discovery is the reliance on reference-based analyses, which is an effective approach if the disease locus and transcript structures in proband and reference assemblies are shared, but if they differ, the methods break down. This confound may be amplified in disorders where reliance on a consensus human reference may be insensitive to cryptic sequences unique to a founder haplotype. Conventional approaches can also be challenged by mechanistic interpretation in late onset disorders, where variants that contribute to risk may exert subtle effects that do not impede normal health and development for a large proportion of the patient’s life.

One illustrative example of such an elusive Mendelian disorder is X-linked Dystonia-Parkinsonism (XDP), an adult-onset neurodegenerative disease that has presented a unique challenge to conventional gene discovery. XDP is endemic to the island of Panay, Philippines, where its prevalence is reported to be 5.74 cases per 100,000 individuals with a mean age of onset of 39.7 years (Lee et al., 2011). The clinical phenotype documented most frequently combines features of dystonia and parkinsonism in a characteristic temporal progression, beginning with hyperkinetic symptoms at early stages and progressing to predominantly hypokinetic movements at later stages (Evidente et al., 2002; Lee et al., 1991; Lee et al., 2002; Lee et al., 2011). Over the past three decades, conventional genetic approaches have been used to map the XDP causal locus on the X chromosome in search of the pathogenic gene lesion (Domingo et al., 2015; Graeber et al., 1992; Graeber and Müller, 1992; Haberhausen et al., 1995; Kupke et al., 1992; Makino et al., 2007; Müller et al., 1994; Németh et al., 1999; Nolte et al., 2003; Wilhelmsen et al., 1991). These studies revealed an apparently identical disease haplotype shared by all probands consisting of seven variants: five single nucleotide variants (SNVs), designated in the literature as <u>D</u>isease-specific <u>S</u>equence <u>C</u>hanges (DSC) − 1,2,3,10,12; a 48-bp deletion; and a 2,627 bp SINE-VNTR-Alu (SVA)-type retrotransposon insertion within a 449 kb region (Figure 1) (Makino et al., 2007). To date there has been no evidence of discriminating alleles between probands and/or recombination events creating partial haplotypes, nor have these variants been detected in ethnically-matched control individuals.

**Figure 1.**
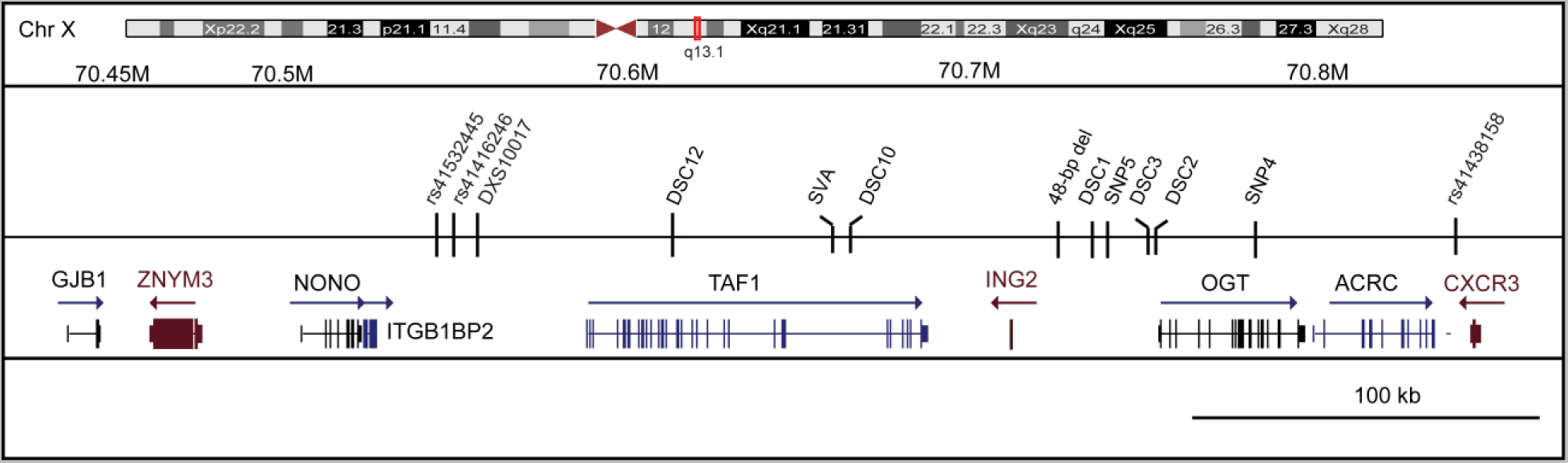
Previously described XDP linkage region. The genomic segment previously associated with XDP on chromosome Xq13.1 is shown. Seven variants have been reported to be shared among XDP probands and not observed in controls: five single nucleotide variants, annotated in the literature as <u>D</u>isease-specific <u>S</u>equence <u>C</u>hanges (DSCs)-1,2,3,10,12; an SVA retrotransposon inserted antisense to TAF1; and a 48-bp deletion. Also shown are flanking markers from a recent genotyping study to narrow the shared region.

The major challenge in understanding the etiology of XDP has been that none of these disease-specific variants have annotated functions. Three (DSC10, 12, and the SVA) fall within introns of the human *TAF1* gene, whereas the remaining four are localized within an intergenic region 3’ to *TAF1* that reportedly contains five unconventional exons, designated as a Multiple Transcript System (MTS), which may be transcribed as distinct RNAs and/or spliced to *TAF1* exons (Herzfeld et al., 2007; Makino et al., 2007; Nolte et al., 2003). These observations have raised the possibility that a defect in *TAF1* may somehow underlie XDP pathogenesis. *TAF1* encodes TATA-Binding Protein (TBP)-Associated Factor-1 (TAF1), a component of the TFIID complex which mediates transcription by RNA polymerase II (Pol II) (Anandapadamanaban et al., 2013; Kandiah et al., 2014; Louder et al., 2016; Thomas and Chiang, 2006) and has emerged in recent years as a significant disease target. In addition to the XDP-related sequence variants in this gene, other studies have reported coding variation in *TAF1* to be associated with severe neurodevelopmental defects and intellectual disability (O’Rawe et al., 2015) as well as various cancers (McKeown et al., 2014; Oh et al., 2017; Zhao et al., 2013). Given the essential function of TAF1 in transcriptional regulation in all cells, it is not known how these reported sequence variants may cause tissue-specific defects and/or highly specific clinical phenotypes.

In this study, we investigated XDP as an exemplar of an unsolved Mendelian disorder arising from a founder haplotype in an isolated population. We hypothesized that the full genetic diversity of XDP may not have been captured by previous genetic approaches, and that unbiased assembly of the genome and transcriptome spanning the XDP haplotype could reveal novel sequences or aberrant transcripts unique to XDP probands. We thus approached this problem by integrating multiple short and long-read sequencing technologies and reference-free assembly approaches in XDP model cell lines. Our results identified novel genomic variants and assembled transcripts that were shared among XDP probands, but not observed in controls, including aberrant splicing and partial retention of intronic sequence in proximity to the disease-specific SVA insertion within intron 32 of *TAF1.* This intron retention (IR) correlated with an overall reduction in expression of the canonical transcript, cTAF1, in XDP probands and a co-expression network strongly enriched for differentially expressed genes (DEGs) associated with neurodevelopmental pathways. Excision of the SVA via CRISPR/Cas9-based genome editing rescued the IR event and normalized *TAF1* expression differences. Collectively these data offer new insight into the transcript structure of *TAF1* in neural cells, implicate a unique genomic cause for XDP, and provide a potential roadmap for integrated, reference-free genome and transcriptome assemblies in population isolates.

## Results

### Establishment of an XDP familial cohort

Samples were obtained from 120 individuals: 57 affected hemizygous males, 23 heterozygous carrier females, 32 unaffected family members, and 8 males who were carriers of the haplotype but had not reached the median age of disease onset and were asymptomatic at the time of sample collection, referred to herein as non-manifesting carriers (NMCs; see Table S1). The cohort included 19 archival specimens which were previously described (Nolte et al., 2003). Pedigrees and clinical characteristics of subjects are summarized in Table S1. Affected males exhibited variable degrees of dystonia and/or parkinsonism, consistent with clinical reports of XDP (Lee et al., 2011), and were positive for the known XDP haplotype markers based on PCR amplification of genomic DNA (gDNA) extracted from blood, followed by Sanger sequencing of amplicons. The heterozygous carrier females were positive for the haplotype but appeared neurologically normal upon exam, as did the haplotype-negative control relatives.

### Genome assembly and deep sequencing of the XDP haplotype reveal novel shared sequences and narrow the causal locus

We hypothesized that the XDP founder haplotype might include sequences unique to the Panay population and cryptic to the current human genome assembly, which could complicate discovery of XDP-specific variants. Because all previous studies of the XDP haplotype have reported seven known variants that were shared by all probands, with no discriminatory alleles among probands, we also explored whether the region may be resistant to recombination due to an existing structural rearrangement. To test these possibilities, we used four strategies: (1) reference-free, *de novo* assembly of the XDP haplotype among probands and control relatives using Illumina paired-end and 10X Genomics linked-read sequencing; (2) Pacific Biosciences long-read single molecule real-time sequencing (PacBio SMRT) of bacterial artificial chromosome (BAC) clones of the XDP haplotype; (3) long-insert “jumping library” whole genome sequencing (liWGS), which we previously developed to probe for structural variation (Collins et al., 2017; Hanscom and Talkowski, 2014); and (4) targeted capture (CapSeq) for dense tiling and deep sequencing of the XDP genomic region in 117 subjects.

For *de novo* assembly, Illumina paired-end 250 base pair (bp) reads were initially assembled from six samples representing two independent XDP families (pedigrees 22 and 27, Table S1) using the algorithms DISCOVAR and DISCOVAR *de novo* (Weisenfeld et al., 2014). These analyses generated a contiguous 410,455 nucleotide haplotype block spanning the complete genomic segment previously linked to the XDP locus, including 2,106 nucleotides not observed in the reference sequence (ChrX:70429400-70841578). We also used PacBio SMRT technology to sequence BAC clones derived from a single proband (pedigree 12) covering a 200 kb segment spanning *TAF1* with the SVA (ChrX:70546230-70747084). The assembled contig (average read length after filtering = 10,416 bp) confirmed all Illumina results and assembled the complete SVA sequence (2,712 bp; Figure S1). The 3.5 kb liWGS was performed on six individuals (four probands and two unaffected relatives; pedigrees 22 and 27, Table S1) using a previously described pipeline to probe structural variation at approximately 5 kb resolution (Collins et al., 2017; Hanscom and Talkowski, 2014). These analyses revealed no structural rearrangements unique to probands that spanned the XDP haplotype, or any other region on the X chromosome, that might inhibit recombination of the founder haplotype.

The *de novo* assembly produced a haplotype region that we densely tiled using 120mer long RNA baits with deep sequencing of the entire segment in 117 individuals (463 kb including flanking regions; Table S1), generating a median of 85X read depth and coverage of 96% of all targeted bases. The CapSeq and WGS assembly revealed considerable diversity of variation spanning the XDP haplotype, including 497 single nucleotide variants (SNVs) and 68 insertion/deletions (indels), 25% of which were novel compared to the 1000 Genomes Project (Genomes Project et al., 2015), Exome Aggregation Consortium (ExAC) (Lek et al., 2016), and dbSNP. We detected all seven previously annotated sequence variants as well as 40 additional variants that segregated with disease status in this cohort for a total of 47 variants associated with the XDP haplotype (38 SNVs, 7 indels, the SVA insertion, and the 48 bp deletion; Figure 2A, Table S2, Figure S2). Novel DSCs are annotated in Figure 2 as DSCn to maintain consistency with the existing XDP literature and in Table S2 with standard nomenclature for integration with human genetic reference maps.

**Figure 2.**
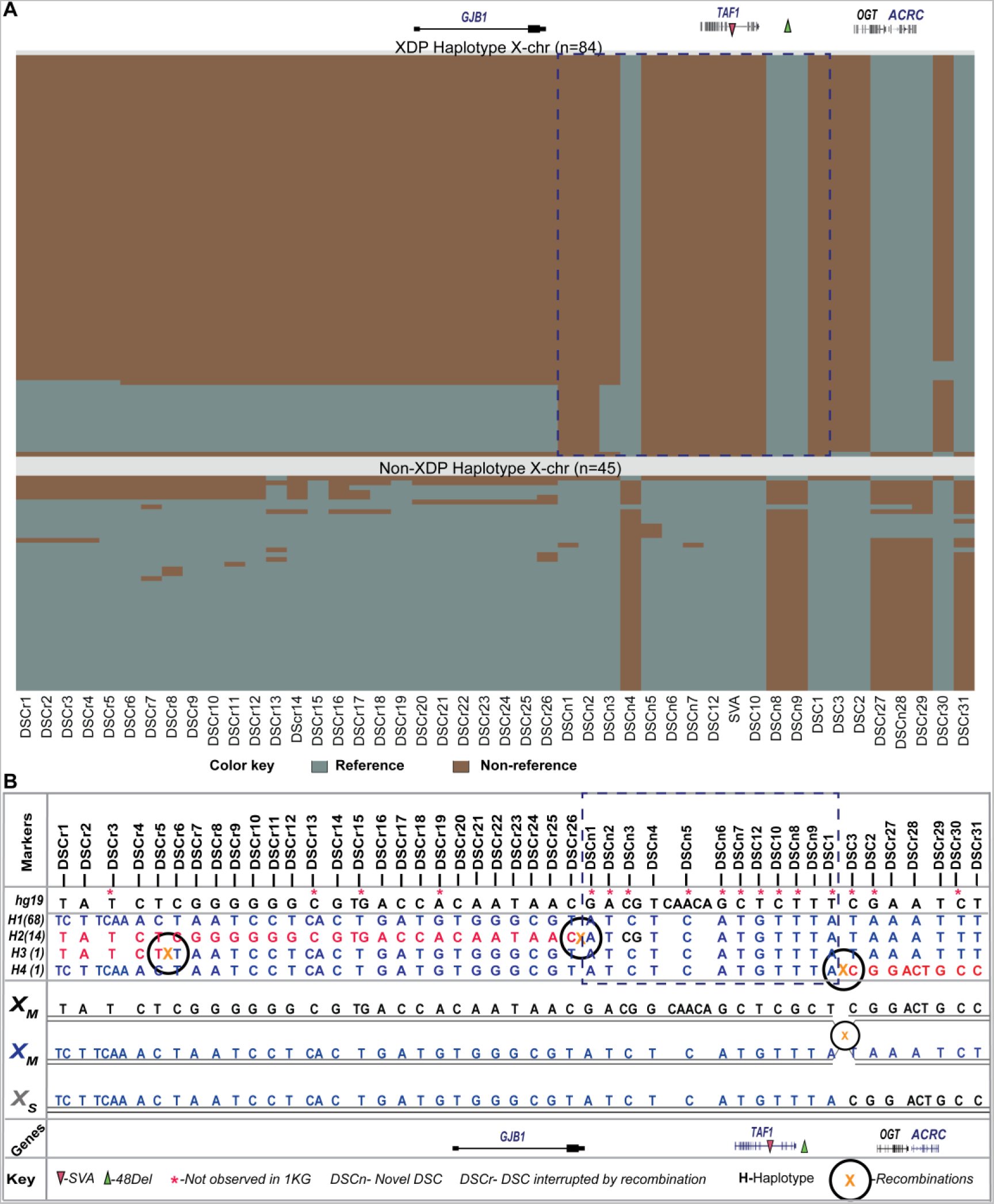
Complete allelic architecture of shared variants among XDP probands. (A) Genome assembly and targeted capture of the XDP haplotype region revealed all previously described variants, 40 novel variants shared among probands, and three recombinations. Data provided for XDP probands and non-manifesting carrier (NMC) males (n = 57 probands + 8 NMC haplotypes), as well as the phased XDP haplotype from carrier females (n = 19). Non-XDP haplotypes are provided for 26 unaffected relatives and the phased control haplotype from carrier females (n = 19 haplotypes). Colors represent reference and non-reference alleles at 47 loci compared to the reference human genome and with differential allelic profiles between probands and unaffected members. (B) Annotations for the 47 XDP-associated loci indicating positions, recombinations, and alleles in the different haplotype groups, as well as the genes containing the specified variations. One recombination was observed within Family 27 to produce haplotype H4, which occurred between DSC1 and DSC3. All alleles for this H4 haplotype are shown with the recombination site: XM = maternal haplotype, XS = Recombinant haplotype. The class of variation and dbSNP ID for the known SNVs and indels are listed in Table S2. A dotted rectangle represents the likely narrowed XDP region shared when considering recombinations, though it includes the reversion to the reference allele observed at DSCn3. 1KG= 1000 genomes; (See also Figure S2).

The assembly and CapSeq data also identified three independent recombinations among XDP probands, the first such recombinations of the XDP haplotype detected to date (two historical recombinations as well as one observed in pedigree 27; Table S1). These recombinations produced four distinct haplotypes, narrowing the shared region among probands to 219.7 kb (Figure 2B). The most common haplotype, H1 (n = 68), consisted of all 47 shared variants and most likely underwent recombination to generate the derivative haplotypes (Figure 2; see Supplemental Information for haplotype details). The second most frequent haplotype involved a recombination proximal to DSCn1 and a reversion to the reference allele at position 70521288 (DSCn3) compared to H1 (H2, n = 14). Among the haplotypes, 13 variants fully segregated with the phenotype in all probands and were not altered by recombination (Figure 2). Variants that defined the H1 haplotype, but no longer segregated with disease due to the observed recombinations, have been annotated as DSCs altered by recombination (DSCr). Sanger validation was performed for selected shared variants predicted from the CapSeq, and 100% of predicted variants were confirmed (Table S2). Collectively, these analyses revealed greater allelic diversity in XDP than had been realized, while further suggesting that all probands share a common core region consisting of at most a 219.7 kb segment (or 203.6 kb if we use the DSCn3 reversion as the proximal flanking point) encompassing *TAF1* exclusively, and most likely reflecting the causal locus.

### Cellular modeling of XDP

To interrogate the transcript structure of this locus and probe for genotypic differences in expression patterns, we established a collection of XDP and control cell lines consisting of: (1) primary fibroblasts from 13 probands, 12 heterozygous female carriers, and 20 unaffected relatives; and (2) iPSCs from 5 XDP probands, 4 female carriers, and 3 unaffected relatives, with 2 clones per individual (24 total clones; Table S1). Pluripotency analysis of XDP and control iPSCs has been previously reported (Ito et al., 2016). Figure S3 depicts similar characterization of iPSCs derived from the female carriers demonstrating expression of standard markers of pluripotency and stemness, trilineage potential, and normal karyotypes. All clones were differentiated in parallel to generate neural stem cells (NSCs) as previously described (Ito et al., 2016) and cortical neurons based on overexpression of neurogenin-2 (NGN2) (Zhang et al., 2013). Expression profiling of the neural derivative cell lines showed the expected segregation of NSC vs. mature neuronal markers in the respective cell types (Figure 3A). Some variability in marker expression was noted across lines, though there were no consistent differences by genotype with the exception of *FOXG1*, which was downregulated in XDP vs. control NSCs (Figure 3A). Although *CUX1* expression has been described in NGN2-induced neurons, in the XDP and control lines it was higher at the NSC stage, which may be consistent with reports that it also marks certain neural progenitors (Nieto et al., 2004). Immunostaining of NGN2-neurons revealed characteristic morphologies with dense networks of processes robustly labeled by doublecortin, MAP2, and Tuj1 (Figure 3B).

**Figure 3.**
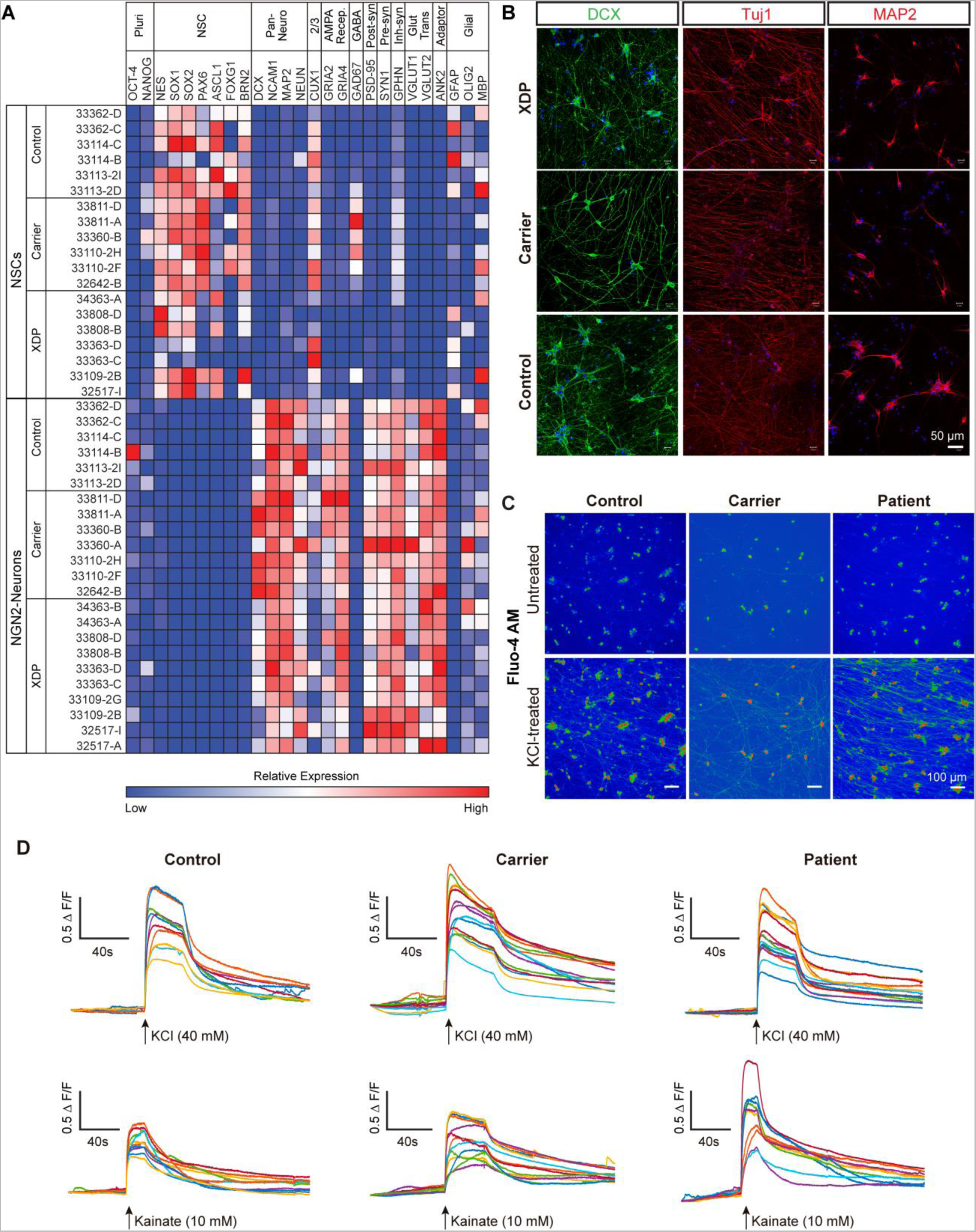
Characterization of iPSC-derived NSCs and NGN2 induced neurons. (A) Heatmap of relative expression of pluripotency, neural stem cell, mature neuronal and glial genes in NSCs and NGN2-induced neurons based on RNAseq reads. (B) Representative images from proband, carrier female, and unaffected control NGN2-induced neurons showing comparable morphologies with dense networks of processes immunopositive for doublecortin (DCX), βIII-tubulin/Tuj, and MAP2. (C) Activity-dependent Ca^2+^ mobilization in NGN2-neurons visualized via the fluorescent indicator Fluo-4AM. Upper and lower panels show raw fluo-4AM fluorescence before and after, respectively, 40 mM KCl treatment in control (left panels), carrier (middle panels) and XDP (right panels) lines. (D) Representative traces show relative change in fluorescence intensity (ΔF/F) upon application of 40 mM KCl (upper panels) and 10 mM kainate (lower panels) in control (left panels), carrier (middle panels) and patient (right panels) lines. Colored traces represent individual cells (n = 10-15 cells).

Functional maturity was evaluated based on activity-dependent intracellular calcium mobilization. Neurons loaded with the calcium indicator dye, Fluo-4, exhibited robust calcium influx elicited by both KCl (40 mM) depolarization and the glutamate receptor agonist, kainate (10 mM) (Figure 3C-D), the latter of which could be blocked by the AMPA/kainate receptor antagonist CNQX (20 μM), demonstrating the specificity of the response.

### XDP cellular models exhibit differential expression of TAF1 transcripts and partial retention of an intronic sequence proximal to the SVA insertion

To evaluate expression changes associated with the XDP haplotype and assemble the complete transcript structure of *TAF1* in all three cell types, we used: (1) strand-specific dUTP-RNAseq and Illumina paired-end sequencing (median = 39.6M paired-reads per clone); and (2) a targeted mRNA capture method we developed using the same baits from the DNA CapSeq (referred to here as RNA CapSeq), which tiled all coding and noncoding transcripts spanning the region. The method successfully yielded an approximately 150-fold increase in coverage of targeted transcripts (median = 2.7M paired-end reads spanning the segment per clone). Given the challenges associated with *de novo* assembly of novel transcripts (Chang et al., 2015), we required support from both methods to identify novel transcripts, which were then manually inspected for all novel junctions. Initial analyses identified *TAF1* as the only differentially expressed gene between XDP probands and unaffected controls spanning the XDP linkage region, further supporting the likelihood that the narrowed segment encompassed the causal locus. Expression of *TAF1* was reduced among XDP probands in both fibroblasts (*p*-value = 2.0 x 10^-5^) and NSCs (*p*-value = 1.2 x 10^-3^), but not in the induced neurons (*p*-value = 0.82), and we thus focused our analyses on this locus.

*De novo* assembly yielded four *TAF1* transcripts previously annotated and five novel isoforms in these cell lines (Figure 4A, Table S3-S4). The canonical *TAF1* transcript, cTAF1, was the predominant species in all cell types, representing 70.6%, 56.6%, and 33.6% of total *TAF1* expression in fibroblasts, NSCs, and neurons, respectively. *TAF1* contains a putative neuron-specific isoform, nTAF1, which differs from cTAF1 by 6 bp derived from an alternative exon annotated as 34’ (Makino et al., 2007). Consistent with previous studies, the canonical nTAF1 isoform was the second predominant transcript in mature neurons (24.2% of total *TAF1* expression), but our analyses revealed that 57.6% of all transcripts expressed in neurons also incorporated exon 34’. The canonical nTAF1 transcript was not expressed in fibroblasts and was only detectable at low levels in less than half of all NSC clones. Across all transcripts, five isoforms shared canonical exons but differed from cTAF1 due to truncation of exon 5 and/or inclusion of alternative exons (Figure 4A, Table S3, see Supplemental Information). Although we observed multiple novel junctions that suggested transcription may extend beyond *TAF1*, we did not detect the previously proposed MTS exons in these cells (Herzfeld et al., 2007; Nolte et al., 2003).

**Figure 4.**
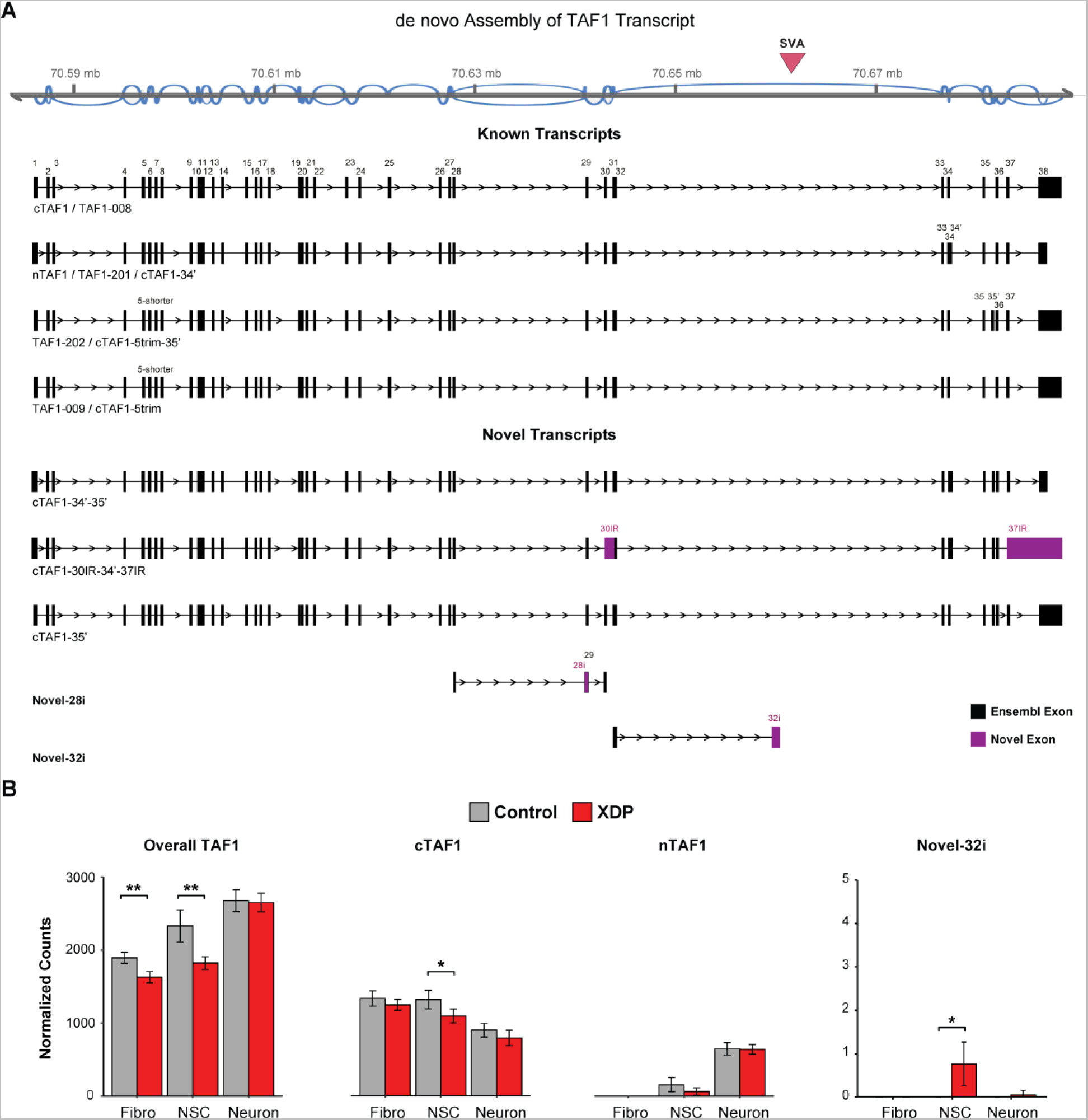
De novo assembly of TAF1 transcript structure and differential expression of splice variants. (A) Sashimi plot (top panel) displays the predominant splicing patterns detected by RNASeq (read depth ≥ 10). Pink triangle indicates position of the XDP-specific SVA insertion. Transcript structure plot derived from de novo assembly depicts TAF1 isoforms previously annotated in Ensembl and novel splice variants detected here. Within each transcript, boxes denote exons either in grey (Ensembl-annotated exon) or purple (novel). Box numbers indicate alternative exons compared to cTAF1/TAF1-008 exons. (B) Barplots demonstrate overall TAF1 gene expression and the expression of three selected isoforms between control (grey) and XDP (red) samples across three cell types. We found that the significant difference in gene expression was driven by cTAF1 and it’s derivative isoforms, while no differences were detected in nTAF1 expression or all transcripts that included splicing of exon 34’ in the induced neurons (Figure S6, Table S4). The error bars reflect ±1.96 fold of S.E.M. “*”: p < 0.05; “**”: p < 0.005.

Comparison of TAF1 transcript levels between genotypes revealed that the overall differential expression reflected reduced expression of cTAF1 and several derivative isoforms in fibroblasts and NSCs, but not neurons (Figure 4B, Table S5-S6; Figure S4-S5; see Supplemental Information). The nTAF1 transcript did not differ betwen XDP and control neurons, the only tissue in which it could be robustly quantified (Table S5). We scrutinized the expression patterns associated with cTAF1 and several related isoforms and found that the overall reduction in expression among XDP fibroblasts and NSCs (Table S6) was largely driven by decreased exon usage toward the 3’ end of the gene, and the pattern was most pronounced for exons distal to the SVA insertion site (Figure S4A-C). We also discovered that the only exception to the otherwise uniformly decreased expression among these transcripts was an assembled transcript that was up-regulated in XDP probands and terminated within intron 32 (annotated here as Novel-32i in Figure 4B and Table S5-S6). In NSCs, the Novel-32i transcript involved splicing to a position in intron 32 just 716 bp proximal to the SVA, which was a junction never detected in control NSCs (Figure 4B and S6A, Table S5). We also discovered a striking intron retention (IR) signature in XDP NSCs and, to a lesser extent, in fibroblasts that was never observed in controls; intronic sequence was transcribed spanning the length of intron 32 and terminating 716 bases prior to the SVA junction site (corresponding to the 3’ segment of the SVA given its antisense insertion relative to *TAF1*; Figure 5A). We did not detect inclusion of any SVA sequence in these transcripts or retention of intron 32 distal to the SVA insertion site.

**Figure 5.**
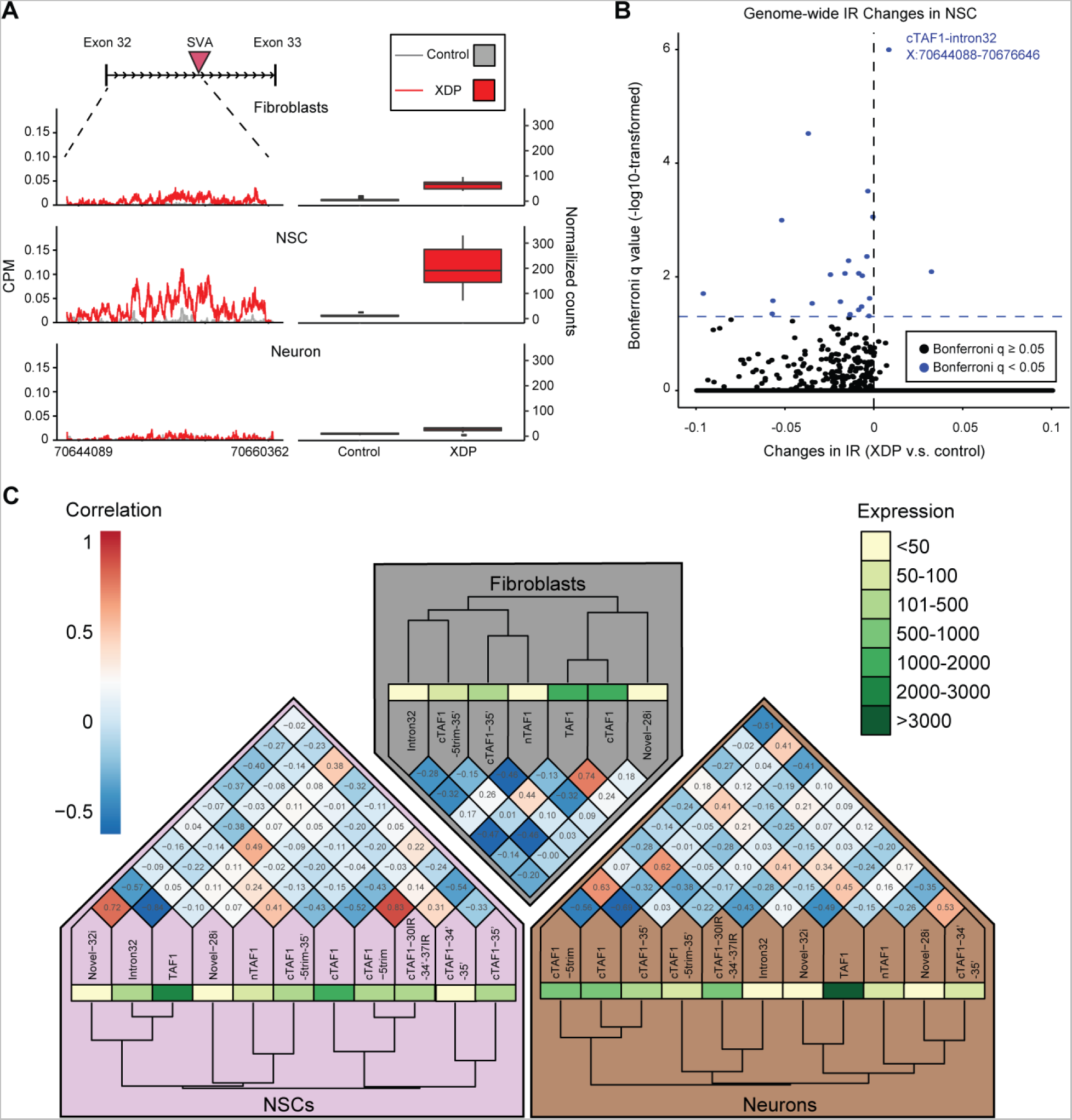
Expression of TAF1 intron 32 and its correlation with TAF1 transcripts. (A) The line plots (left) show the detectable expression by nucleotide position (CPM) of control (grey) and XDP (red) samples from the first base of TAF1 intron 32 to the insertion point of the SVA in three cell types. The boxplots (right) demonstrate the overall expression (normalized counts) and 1.5 fold of interquartile range of the intron retention (IR) event in the same region for control (grey) and XDP (red) samples. (B) The volcano plot displays genome-wide levels of IR among all 258,852 annotated introns among control and XDP samples in NSCs (x-axis) plotted against their significance levels (y-axis, log10 transformed). Those introns with significant (Bonferroni p-value < 0.05) IR changes are marked in blue. (C) The correlation between the expression levels of all TAF1 transcripts is displayed for each tissue with cluster dendrograms displaying the relationships between transcripts. The colored bar on the right provides the expression levels of each transcript, and the colored grid on the left displays the corresponding strength of correlation, with deeper colors representing stronger positive (red) or negative (blue) correlations. These data show that the intron 32 IR events are positively correlated with each other, and negatively correlated with overall TAF1 expression.

The observed IR patterns were associated with significant differences in cTAF expression between XDP and control cells, with the strongest IR expression peak in NSCs representing ~12% of the average depth of cTAF1 exon expression (Figure 5A and S5A). The aberrant splicing and IR were not observed in XDP neurons, nor has the pattern ever been observed in studies of comparable neural progenitor cells from our group (Sugathan et al., 2014). To empirically confirm that this IR event was an unusual expression pattern relative to null expectations, we compared genome-wide IR patterns between XDP and controls. Across the entire transcriptome, we surveyed 258,852 annotated introns and observed differential retention of 24 introns, of which the *TAF1* intron 32 IR was the most statistically robust, irrespective of directionality, and achieved genome-wide significance (corrected *p*-value = 1 x 10^-6^; Figure 5B and Figure S6D). Notably, evaluation of the correlation structure among all *TAF1* assembled transcripts revealed the intron 32 IR event to be significantly negatively correlated to cTAF1 expression in NSCs, and to overall *TAF1* expression in both NSCs and fibroblasts (Figure 5C), suggesting that retention of this intronic sequence in proximity to the SVA insertion could be driving the reduction in overall *TAF1* transcription in XDP.

### CRISPR/Cas9-mediated excision of the SVA abolishes the intron 32 retention in XDP cells

Given the proximity of the retained intron 32 sequence to the SVA, we hypothesized that the retrotransposon insertion interfered with splicing in this region to produce aberrant transcripts. We evaluated this possibility using CRISPR/Cas9-mediated gene editing to ablate the SVA from an XDP iPSC clone using a dual-guide RNA approach that targeted sequences 5’ and 3’ to the SVA insertion site (Hendriks et al., 2015). Eleven clones were generated in which the 2.6 kb SVA was completely removed. Two clones had the same precise deletion points, which excised the entire SVA as well as 53 nucleotides, representing the sequence between the SVA and the flanking protospacer adjacent motif (PAM) sites (Figure S7A-B). No additional sequence changes or indels were observed following non-homologous end-joining-mediated repair of the 5’ and 3’ cut ends (Figure S7A). These clones were expanded, confirmed to retain normal karyotypes and expression of pluripotency markers, and differentiated to NSCs and NGN2-induced neurons (Figure S7C-F). In NSCs, excision of the SVA completely rescued the intron 32 IR event and aberrant splicing, as the Novel-32i transcript was no longer detectable. Remarkably, removal of the SVA also normalized overall *TAF1* expression, as levels in the edited clones were indistinguishable from that in controls (p-value = 0.8, Figure 6).

**Figure 6.**
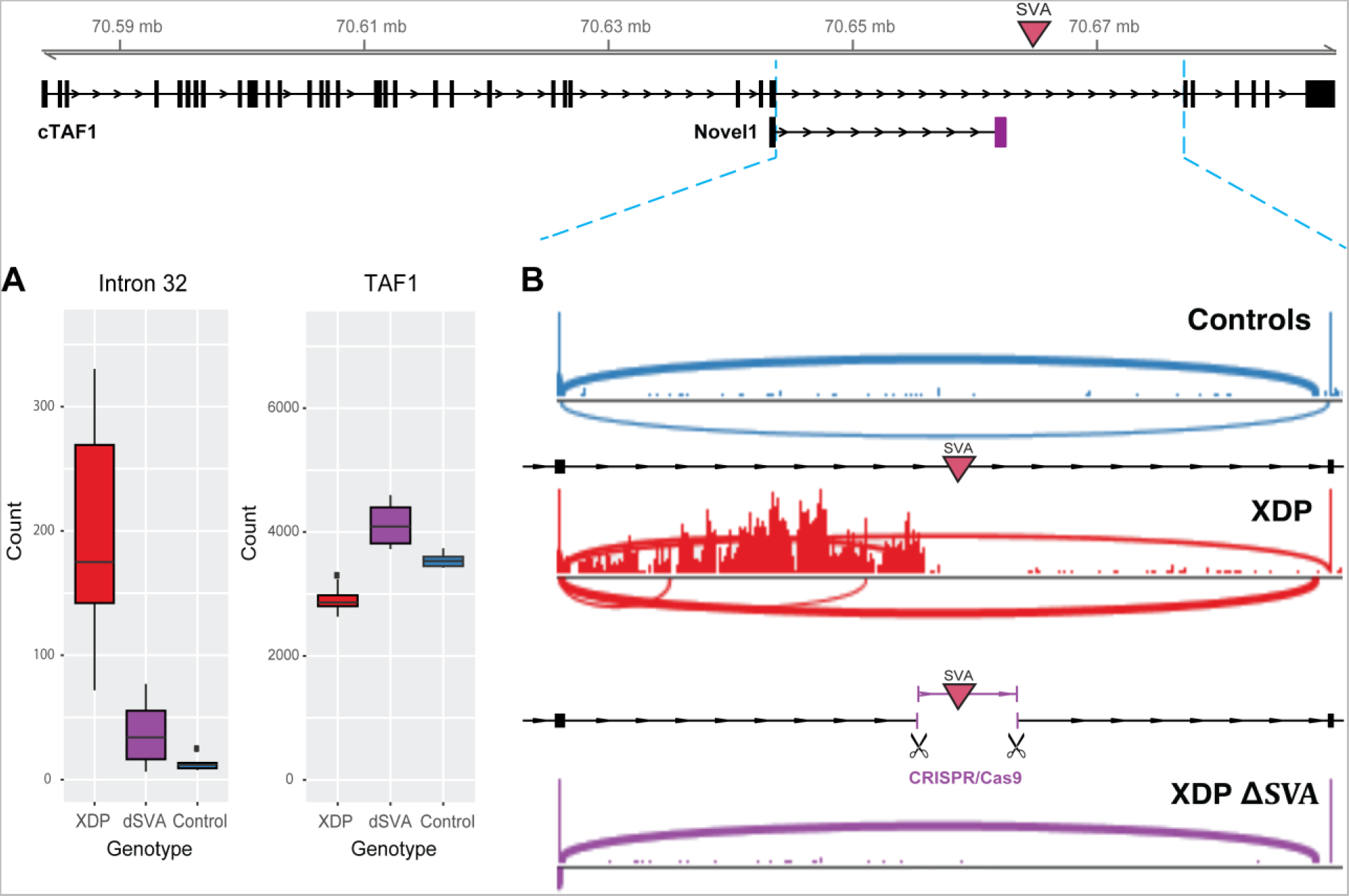
Excision of the SVA rescues aberrant splicing and expression in intron 32 and expression of TAF1. (A) Boxplots of overall expression of intron 32 (left) and TAF1 (right) as normalized counts. (B) Sashimi plots of RNA-seq data in NSCs from probands, controls and CRISPR-modified NSCs from probands with excised SVA (ΔSVA) shows normalization of the TAF1 intron 32 IR event and rescue of overall TAF1 expression to normal levels observed in controls following excision of the SVA.

### Transcriptome-wide molecular signatures reveal XDP-specific alterations in networks related to neurodevelopment and other dystonia genes

To further interrogate the potential downstream functional consequences of XDP-related sequence variation, we characterized global gene expression signatures in all cell lines with functional enrichment analysis (gene ontologies and pathways) of DEGs and weighted gene co-expression network analysis (WGCNA) on all samples. Given the complexity of these analyses, the limited power from a relatively small number of available iPSC-derivative lines, and the potential confounds of inter-iPSC clone variability, we used principal component analysis (PCA) to identify clusters of samples with similar profiles, not influenced by genotype, but contributing to expression heterogeneity. PCA factor adjustment was incorporated into the gene count modeling and statistical analyses. Both unadjusted and adjusted *p*-values are reported, but genes were only considered to be differentially expressed at *p-*value < 0.1 after adjustment for multiple comparisons. Fold changes, associated probabilities, and adjusted *p*-values at multiple significance thresholds for all genes are provided in Table S7.

By these criteria, the number of DEGs and magnitude of expression changes were modest and comparable by cell type (fibroblasts, 37; NSCs, 65; neurons, 56). However, the transcriptional profiles did reveal correlations with similar Gene Ontology (GO) terms, modules, and co-expression networks across multiple cell types. Two genes were significantly altered in multiple cell types: *TAF1* (down-regulated in XDP fibroblasts and NSCs), and *SNHG17*, a noncoding RNA that was downregulated in XDP NSCs and neurons. In fibroblasts, the most significant GO term enriched in DEGs (cerebellar granular layer morphogenesis; *p*-value = 2 x 10^-6^) was represented by three genes, *NRXN1*, *CBLN1* and *GRID2*, all of which have been associated with both neurodevelopmental and neurodegenerative disorders (Greenwood et al., 2016; Kalkan et al., 2016; Kim et al., 2008; Krishnan et al., 2017). The co-expression network with strongest DEG enrichment in fibroblasts (Module 8; enrichment *p*-value = 7 x 10^-3^) was associated with the GO term “synaptic transmission” (*p*-value = 1.9 x 10^-12^) and enriched for genes linked to other movement disorders (*HPCA*, *ADCY5, GNAO1, TUBB4A, TUBA4A, LINGO1, LINGO2*; *p*-value = 2 x 10^-4^). This same term, “synaptic transmission”, was also associated with the module most enriched for DEGs in NSCs (*p*-value = 2.6 x 10^-10^). The enrichment of DEGs within this module was highly significant (*p*-value = 4.5 x 10^-8^), as 16.7% of genes were significant at adjusted *p*-value < 0.1, and all genes were differentially expressed at nominal p < 0.15. Protein-protein interaction network analysis also revealed strong enrichment for physical interactions among these loci (*p*-value = 5.64 x 10^-11^), and this network involved a substantial number of genes previously associated with neurodevelopmental disorders, including *TBR1* (Notwell et al., 2016), *FOXG1* (Mariani et al., 2015; Won et al., 2016), and *GRM5* (Guo et al., 2016) (Figure 7). Consistent with this pattern, the top DEG in neurons, *VAMP1*, encodes part of the synaptic vesicle machinery (Bourassa et al., 2012) and exhibited the greatest genotypic difference in expression level of any marker gene among the three cell types. These data suggest that XDP is associated with altered expression of genes associated with neurodevelopmental processes, either as a consequence of reduced of *TAF1* expression or of the overall disease process.

**Figure 7.**
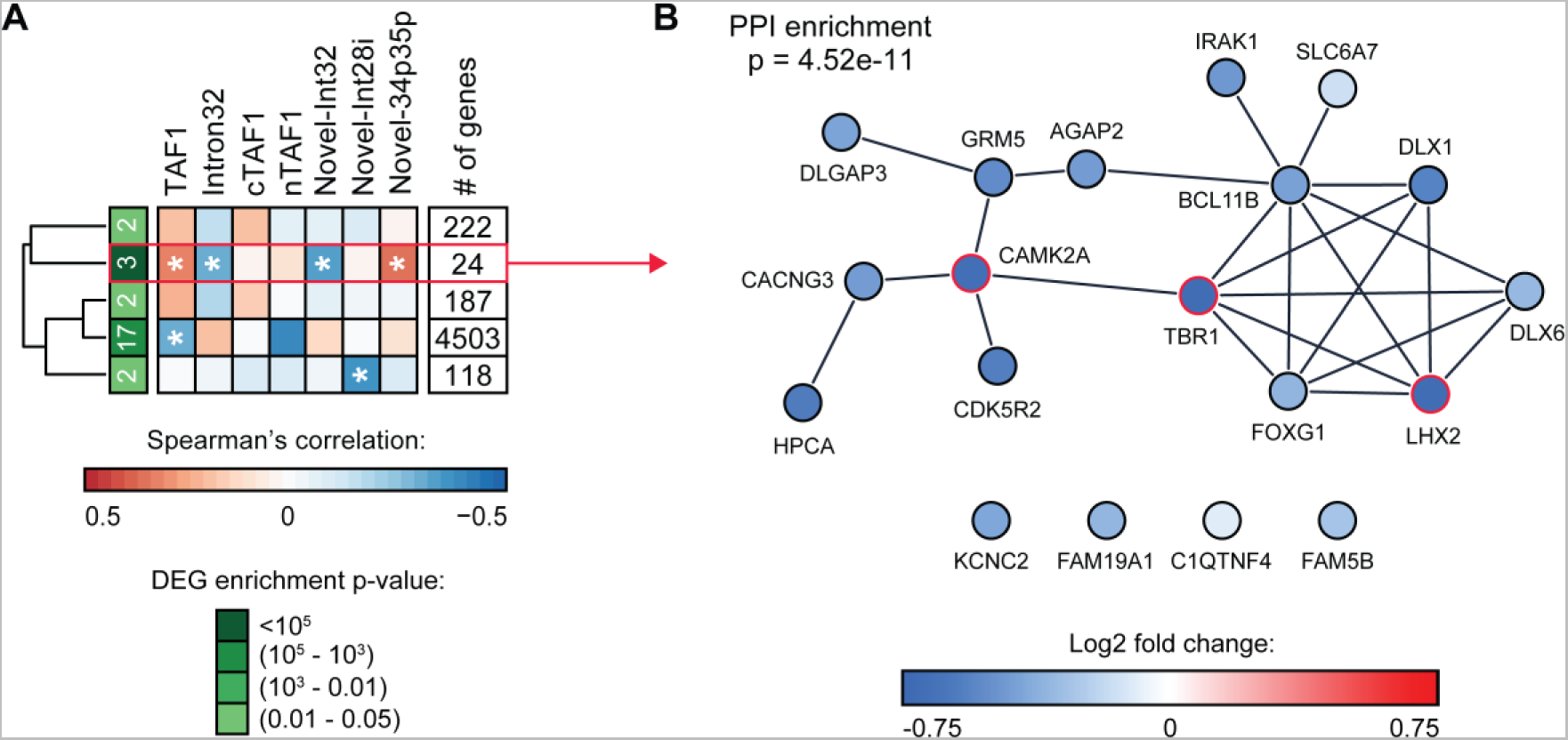
Strongest co-expression modules identified in NSCs are enriched for differentially expressed genes and neurodevelopmental loci. (A) Modules with significant enrichment for differentially expressed genes (DEGs) and the correlation of these modules to expression of TAF1, intron 32 and multiple TAF1 transcripts are shown. * indicates significant correlation with p-value < 0.05. The module with strongest enrichment p-value is highlighted in purple. (B) A protein-protein interaction network built from the module with the strongest enrichment of DEGs (module 24) using string-db. The color of the nodes indicates log2 fold change between probands and controls based on differential expression analysis. Red nodes indicate statistically significant DEGs.

## Discussion

The impact of rare noncoding regulatory variation in human disease represents a substantial knowledge void and is currently an area of intensive study. There are relatively few examples of noncoding regulatory variants causally linked to Mendelian disorders, yet it is known that some dominant-acting noncoding mutations can confer substantial risk (Mathelier et. al, 2015). The relative contribution of retrotransposition to disease represents another area of uncertainty, as such events occur relatively frequently in the germline and somatic tissues. In the human genome there are three classes of active retroelements (LINE or L1, ALU, SVA) which are currently linked to 85 human diseases. Although some of these diseases involve insertions within exons, many are associated with intronic insertions which affect transcription and splicing (Kaer and Speek, 2013). Consistent with that pattern, the genome and transcriptome assembly reported here narrowed the causal locus in XDP to a genomic segment including only *TAF1* and discovered that an intronic SVA insertion is associated with altered splicing and expression of the host gene, further supporting the hypothesis that transcriptional interference may be a common consequence of intronic retroelements.

The *de novo* transcriptome assembly, coupled with deep targeted sequencing, enabled an unbiased evaluation of all transcribed sequences spanning the XDP linkage region and identified a truncated *TAF1* transcript that involved the striking transcriptional signature of aberrant splicing and IR associated with XDP. Removal of the SVA rescued this XDP signature, suggesting that the SVA insertion was the predominant driver of the aberrant splicing, IR, and overall decrease in *TAF1* expression in XDP. Alternative splicing events may be classified as exon skipping or inclusion, alternative splice site selection, and IR (Jacob and Smith, 2017; Vaquero-Garcia et al., 2016; Wang and Burge, 2008; Yap and Makeyev, 2016). In each scenario the generation of alternate transcripts is determined by *cis*-acting sequence elements, such as constitutive splice motifs and splicing enhancers/silencers, and *trans*-factors including the core splicing machinery and additional RNA-binding proteins (Wang and Burge, 2008). Until recently, IR events have been regarded as rare consequences of aberrant splicing (Jaillon et al., 2008; Roy and Irimia, 2008), but increasing evidence suggests that they are prevalent throughout mammalian transcriptomes (Braunschweig et al., 2014; Middleton et al., 2017) and constitute a critical mechanism regulating gene expression (Jacob and Smith, 2017; Mauger et al., 2016; Wong et al., 2016; Yap and Makeyev, 2013). This regulation may “fine tune” overall transcript levels, as IR-transcripts typically trigger various quality control mechanisms, including nuclear restriction, nonsense-mediated mRNA decay (NMD), and turnover via exosomes, which prevent their translation (Ge and Porse, 2014; Jacob and Smith, 2017; Yap et al., 2012). As a result, IR-transcripts often undergo rapid turnover, exist at low steady-state levels, and correlate with decreased overall transcript levels, consistent with the pattern we detected in XDP cells.

Based on the normalization of the aberrant splicing and restoration of normal *TAF1* expression following excision of the SVA, we speculate that the retrotransposon insertion in intron 32 of *TAF1* has modified the splicing process in this region. SVAs are the youngest family of retroelements with at least 2700 known insertions and a retrotransposition rate of approximately 1 in 916 births (Wang et al., 2005; Xing et al., 2009). They frequently transduce both 5’ and 3’ flanking sequences and contain multiple 5’ and 3’ splice sites (Damert et al., 2009; Hancks et al., 2009; Kaer and Speek, 2013). Of the 85 known diseases associated with active retroelements, seven (including XDP) are linked to SVAs, five of which involve insertions into introns (Kaer and Speek, 2013), which primarily result in exon skipping and/or exonization of SVA sequences due to activation of cryptic splice sites (Akman et al., 2010; Conley et al., 2005; Hassoun et al., 1994; Kobayashi et al., 1998; Taniguchi-Ikeda et al., 2011; van der Klift et al., 2012; Wilund et al., 2002). All IR detected in intron 32 terminated proximal to the SVA insertion site. A similar pattern was recently reported for an SVA insertion in intron 8 of *CASP8* which resulted in significant IR and decreased exon expression (Stacey et al., 2016). Because intron excision occurs during transcription (Ameur et al., 2011; Khodor et al., 2011; Pandya-Jones and Black, 2009), its precision can vary with changes in the elongation rate of RNA Pol II (Braberg et al., 2013; Fong et al., 2014; Jimeno-Gonzalez et al., 2015). Intronic retroelements may interfere with RNA Pol II elongation in multiple ways, including (1) binding of a competing Pol II and/or other factors which inhibit progression of the Pol II complex transcribing the host gene; (2) changes to local chromatin; and (3) the presence of repetitive guanine-rich motifs which form inhibitory quadruplex structures (Kaer and Speek, 2012; Kejnovsky and Lexa, 2014; Lexa et al., 2014; Savage et al., 2013). These interactions are all examples of transcriptional interference (TI) in *cis* in which the Pol II complex transcribing the host gene may be displaced by some obstacle (Hao et al., 2014; Hao et al., 2017; Shearwin et al., 2005). In this study, the decreased exon usage downstream of exon 32 appears consistent with TI induced by the SVA, and further characterization of the specific pathogenic mechanism will be required in future studies.

These data suggest that XDP may join a growing list of human diseases involving defective RNA splicing, IR, and transposable element insertions (Kaer and Speek, 2013; Scotti and Swanson, 2016; Ward and Cooper, 2010). Extensive studies are required to understand the precise mechanism by which the SVA insertion is driving the aberrant expression signatures observed, but it is encouraging that, for some of these conditions, considerable progress has been made in designing strategies to correct splicing events using small molecules and antisense oligonucleotides (Faravelli et al., 2015; Shimizu-Motohashi et al., 2016; Singh et al., 2015). The potential to normalize this IR event by manipulating the SVA insertion in this study, coupled with rapid advances in genome editing technologies, raises the provocative possibility that *in vivo* manipulation of this sequence could eventually have clinical benefit.

## Author contributions

Conceptualization, M.E.T., D.C.B., X.O.B., L.J.O.; Methodology, T.A., W.T.H, R.Y., D.S., D.G., A.S., B.C., H.B., C.A., A.R., N.I., N.W., D.J., M.E.T., D.C.B.; Formal Analysis; T.A., R.Y., D.G., R.L.C., A.S., B.C., H.B., C.H., C.A., A.R., M.E.T.; Investigation, T.A., W.T.H., R.Y., D.S., D.G., C.A.V., R.L.C., A.S., B.C., H.B., C.H., C.A., J.D.; Resources, C.G., M.D., P.A., N.S.; Writing, T.A., W.T.H., R.Y., D.G., M.E.T., D.C.B.; Supervision, M.E.T., D.C.B., L.J.O. X.O.B.; Funding Acquisition, D.C.B., M.E.T., N.S., L.J.O, X.O.B.

## Acknowledgements

Flow cytometry and sorting services were supported by shared instrumentation grants 1S10OD012027-01A1, 1S10OD016372-01, 1S10RR020936-01, and 1S10RR023440-01A1 at the MGH Department of Pathology Flow and Image Cytometry Research Core. Funding for this study was provided by the MGH Collaborative Center for X-Linked Dystonia-Parkinsonism (D.C.B., M.E.T., N.S.), and National Institutes of Health grants 5P01NS087997 (D.C.B., N.S., L.J.O., X.O.B.), R00MH095867 (M.E.T.), and R01NS102423 (M.E.T. and D.C.B.). Dr. Talkowski is also supported as the Desmond and Ann Heathwood Research Scholar.

## STAR Methods

All critical reagents, cell lines, and software used and/or generated in this study are listed in the Key Resource Table along with corresponding vendor information and/or citations, where appropriate.

### Contact for Reagent and Resource Sharing

All data for this study will be made available in dbGAP (accession number pending). Fibroblast lines have been deposited at the NINDS Human Cell and Data Repository (http://ninds.genetics.org; Rutgers, NJ), and iPSC clones are publicly available from WiCell (www.wicell.org). Further information on cellular resources and reagents should be directed to Dr. D.C. Bragg, and information on genomics datasets should be addressed to Dr. M.E. Talkowski.

### Clinical Evaluation of Subjects and Sample Collection

Subjects recruited for this study included individuals who (1) had a confirmed diagnosis of XDP based on prior genetic testing; (2) exhibited clinical features consistent with XDP and reported ancestry to Panay; and (3) were first-degree relatives of individuals with a confirmed or suspected diagnosis of XDP. Participants were evaluated at Massachusetts General Hospital (Boston, MA) and in regional clinics in Panay Island affiliated with Jose Reyes Memorial Medical Center (Manila, Philippines). The study was approved by institutional review boards at both institutions, and all participants provided written informed consent. Clinical evaluation consisted of videotaped neurological exams with scores recorded for standard rating scales: Burke-Fahn Marsden, Toronto Western Spasmodic Torticollis Rating, and Voice Disability Index (Albanese et al., 2013). In addition, archival DNA specimens from XDP subjects were provided by Dr. Ulrich Müller (University of Giessen; Giessen, Germany). Collection of these samples and clinical evaluation of donor subjects were previously reported (Lee et al., 1991; Nolte et al., 2003). Skin biopsies and fibroblast derivation were performed as previously described (Ito et al., 2016), and fibroblasts were cultured in Dulbecco’s Modified Eagle Medium (DMEM) supplemented with 10% fetal bovine serum (FBS) and 1X penicillin/streptomycin in a humidified incubator at 5% CO_2_.

### PCR-free and Linked-read Deep Whole-genome Sequencing and Assembly

DNA was extracted from blood samples taken from two probands, two female carriers and two controls (pedigrees 12 and 27). Libraries were prepared for paired-end 250 cycle sequencing using standardized PCR-free library production and sequencing on an Illumina HiSeq 2500 at the Broad Institute Genomics Platform. All data was aligned to the human reference genome GRCh38 for reference guided assembly, and reference free assembly was performed using DISCOVAR *de novo*. All methods for data production and assembly are provided in complete detail in Weisenfeld et al. (Weisenfeld et al., 2014). The known XDP-specific variants were verified in the assembled region from the pedigree 12 proband. For haplotype phasing and structural variant detection, three members of pedigree 12 were sequenced using linked-read whole-genome sequencing (10X Genomics, Pleasanton, CA). The library preparation was done by 10X Genomics directly from frozen lymphoblastoid cell lines, and the resulting libraries were sequenced at The Broad Institute on an Illumina HiSeq 2500 platform. Sequenced reads were assembled using the 10X assembler Supernova (http://biorxiv.org/content/early/2016/08/19/070425) and variants were called using GATK (Zheng et al., 2016).

### BAC Generation, Screening, Sequencing, and Assembly

Genomic DNA from proband 33109 (pedigree 12) carrying the entire haplotype region was digested with BamHI and used to generate a BAC library (Amplicon Express; Pullman, WA) in vector pBACe3.6 (Frengen et al., 1999). Nine potentially positive clones were isolated by hybridization using an 856 bp probe located downstream of the SVA (chrX:70,674,877-70,675,733) between exons 34 and 35 of *TAF1*. Genotyping of the DSCs and the SVA was performed as described above and verified one positive clone. A second round of hybridization was carried out to obtain BAC clones containing the 5’ end of *TAF1* using a 201 bp probe located upstream of the SVA (chrX:70,613,608-70,613,809) between exons 21 and 22. Three clones were identified but only one could be verified by PCR. BAC end sequencing using vector primers from the two verified clones suggested they spanned *TAF1* from a region 40 kb upstream of the 5’UTR to 55kb downstream of the 3’ UTR. These two clones were then subjected to long read sequencing (Korlach et al., 2010) to verify their sequence and assemble a contig. A library was made from the BACs (Amplicon Express) using the PacBio 20 Kb library preparation and sequenced on the PacBio RSII instrument (DNA Link, San Diego, CA). A single cell was used to generate 150,292 reads, at an average read length of 5,028. Reads were filtered for vector contamination and quality score, after which 51,210 reads with average length of 10,416 were used to assemble the region. The region was assembled through SMRT-portal (https://github.com/PacificBiosciences/SMRT-Analysis), using HGAP2 protocol with default parameters yet with genome size set to 200 kb. After the assembly, a 201,921bp long single contig was obtained corresponding to hg19 positions 70546230bp to 70747084bp on the X chromosome.

### Jumping Library Preparation and Analysis

Custom ‘jumping libraries’ were prepared as previously described, optimized to 3.5 kb inserts (Hanscom and Talkowski, 2014; Talkowski et al., 2011) and sequenced on an Illumina 2000 platform with 2x51 bp reads. Library barcodes were de-multiplexed and filtered as recommended. Read quality was assessed with FastQC v0.11.2. Reads were trimmed and processed according to the ‘jump shear’ protocol previously described (Redin et al., 2017; Talkowski et al., 2012) and were aligned with BWA-backtrack v0.7.10-r789 (Li and Durbin, 2009) to the human reference genome (GRCh37). Duplicates were marked with SAMBLASTER v0.1.1 (Faust and Hall, 2014). Aligned reads were further processed with with sambamba v0.4.6 (Tarasov et al., 2015). Alignment quality was assessed using PicardTools v1.115, Samtools v1.0, and BamTools v2.2.2 (Li et al., 2009) (Barnett et al., 2011). Structural variants were detected using our published pipeline, Holmes (Brand et al., 2015; Collins et al., 2017).

### DNA Capture-Sequencing (CapSeq) Assay

A capture region on the X chromosome spanning *OGT* to *CXCR3* was targeted using Agilent SureSelect XT design and following the Manufacturer’s instructions (Agilent). Capture libraries were prepared from DNA extracted from proband, carrier and unaffected samples. Three micrograms of gDNA was sheared to approximately 175 bp fragments using the Covaris Focused-ultrasonicator. DNA fragments were end-repaired, adenylated, ligated to adapter oligos and then amplified with 5 cycles of PCR as recommended. After quantification, 750 ng of each amplified DNA sample was hybridized overnight with the capture library. Following capture cleanup, each gDNA library was amplified with an additional 16 cycles of PCR, which also tagged each sample with an index-specific barcode. Final products were quantified using the 2200 TapeStation (Agilent) and pooled for rapid mode sequencing on the Illumina system. Capture libraries were sequenced as 101 bp paired-end reads on an Illumina HiSeq2000 platform

Read pairs were aligned to GRCh37 with BWA-MEM 0.7.5a-r418 (Li H. 2013). Picard Tools and samtools were used to sort and index the alignments and mark duplicates. The CapSeq data resulted in 2.1M reads per sample (read counts range 0.9M - 4.2M per sample) (Table S2). The sequencing data covered 96% of bases of the targeted region with a median coverage of 83X. The Genome Analysis Toolkit (GATK) Haplotype caller v3.5 was used for base quality recalibration, indel realignment, single nucleotide variant (SNV) and indel calling, and genotyping as per published best practice protocols. Because the region of interest is on the X chromosome, the haploidy option was used in GATK (1 for males, 2 for females). SNVs and indels were annotated using Ensembl Variant Effect Predictor (VEP) tool and Scalpel (Fang et al., 2016). GATK haplotype caller detects or reports variants based on alignments, in our case BWA-mem, while Scalpel performs localized micro-assembly of specific regions of interest with the goal of detecting mutations with high accuracy and increased power. One sample (33110, carrier female) had low coverage and thus was excluded from the further analysis of haplotype construction. The females X-chromosomes were phased based on the genotypes in male members of the family whenever possible. Out of 23 female carriers, 19 had a male member to phase the XDP associated region, while three X chromosomes could be phased from the nine control females without a copy of the XDP associated X chromosome.

### Validation to Evaluate Novel DSCs

Validation experiments were performed to estimate specificity for a subset of DSCs detected in this experiment. These analyses were performed for 10 novel variants with sub-threshold sequencing depth across all samples from CapSeq, with all variants and data provided in Table S2. Primers for both PCR and Sanger sequencing were designed to +/-500 bp flanking regions of each variant using Primer3 (v. 4.0). If flanking regions were dominated by low-complexity elements (e.g., LINE, SINE, AT-rich, simple repeats) a nested PCR strategy was employed by designing an outer set of PCR primers, flanking +/-5 kb of variant, in conjunction with inner PCR primers flanking +/-500 bp of variant. PCR was performed using Phusion^®^ High-Fidelity Polymerase, and bands of appropriate size were gel extracted using QIAquick Gel Extraction Kit according to manufacturer’s instructions. Purified PCR products were Sanger sequenced, and results were analyzed using sangeranalyseR and sangerseqR packages in R (Hill et al., 2014), to find the polymorphic sites and heterozygous sites for the carriers. Confirmation rates for detected variants was 100%.

### Generation and Characterization of Induced Pluripotent Stem Cell Lines

The iPSC clones derived from XDP hemizygous males and unaffected control relatives have been previously described, including fibroblast reprogramming, pluripotency evaluation, and trilineage potential (Ito et al., 2016). For this study we similarly reprogrammed fibroblasts from heterozygous XDP female carriers from the same pedigrees using the Cytotune 2.0 Sendai Reprogramming Kit as recommended. Fibroblasts were transduced with Sendai viruses encoding Oct4, Klf4, Sox2 and c-Myc at approximate multiplicities of infection = 3. Cells were fed every other day until day 7, at which point they were replated onto 0.1% gelatin-coated 10 cm plates containing mouse embryonic fibroblasts (MEFs) and switched to hESC medium (7 μL/liter β-mercaptoethanol, 20% Knockout Serum Replacement [KOSR], 2% L-glutamine, 1% Non-essential amino acids [NEAA] and 10 ng/mL basic fibroblast growth factor [bFGF] in DMEM/F12). Colonies were picked by manual dissection, transferred to fresh MEFs, and expanded using mechanical and enzymatic passaging. After initial banking, iPSC colonies were adapted to feeder-free propagation on Geltrex™-coated plates in mTeSR1 medium. Karyotyping was performed by WiCell (Madison, WI) and revealed no chromosomal abnormalities.

Pluripotency of iPSC clones was confirmed based on RT-qPCR and immunostaining for standard markers. For RT-qPCR, RNA was isolated from iPSCs using Zymo DirectZol^®^ RNA miniprep kit, reverse transcribed to cDNA using High Capacity cDNA Reverse Transcription kit, and amplified with primers against *Nanog*, *Oct4*, *Sox2*, *Rex1*, and *Dnmt3B* using PowerUp™ SYBR Green Master Mix, all as recommended. For immunofluorescence, iPSCs were fixed in 4% paraformaldehyde in phosphate-buffered saline (PBS) for 20 minutes at room temperature, washed 3 times in PBS/0.05% Tween 20, permeabilized in PBS/0.1% Triton X-100 for 15 minutes, and then blocked in 4% donkey serum/PBS for 1 hour at room temperature. Cultures were then incubated overnight in primary antibodies against Oct3/4 (1:200 in blocking buffer), Nanog (1:50), SSEA4 (1:200), or Tra-1-60 (1:200). The next day cells were washed 3 times in PBS/0.05% Tween 20), incubated for 1 hr in Alexa Fluor^®^-conjugated secondary antibodies (1:1000 in PBS), washed again in PBS/0.05% Tween 20, and then counterstained with DAPI to visualize nuclei. Images were acquired on a Nikon Eclipse TE2000-U microscope with 20x magnification.

To quantify trilineage potential of iPSC clones, iPSCs were allowed to form embryoid bodies bearing cells of all three germ layers. Cells were incubated in Accutase (1:3 dilution in PBS) for 3 min at 37°C, washed in PBS, and then switched to EB medium (DMEM + 10% KOSR + 1% Pen/Strep) and scraped with trituration to generate small clumps. Clumps were allowed to settle for 5-10 minutes, after which medium was aspirated and cells were gently resuspended in EB medium with the ROCK inhibitor, Y-27632 (4 μM). Cells were seeded onto ultra-low attachment plates to promote EB formation. After 24 hrs, cell suspensions were collected into 15-ml tubes, EBs were allowed to settle, and medium was exchanged to remove ROCK inhibitor before replating. The process was repeated every other day until day 7, at which point they were collected and seeded onto gelatin-coated plates in DMEM +10% FBS + 1% penicillin/streptomycin. Media was exchanged every 3 days until day 14 after initial plating. EBs were then collected, with RNA isolation and cDNA generation performed as described above. Expression of germ layer markers was quantified using Taqman^®^ hPSC ScoreCard™ Panel as recommended. In parallel, some iPSC clones were also evaluated by teratoma formation in mice. For these lines, approximately 1 x 10^6^ cells in Matrigel/PBS were implanted transcutaneously at multiple sites in Fox Chase SCID^®^ mice by the Harvard Genome Modification Facility. Mice were euthanized after 6-8 weeks, and tumors were collected, paraffin embedded, sectioned, and stained with hematoxylin/eosin to identify cells of the three germ layers based on morphology. All research procedures involving animals were approved by the Institutional Animal Care and Use Committee (IACUC) at Harvard University.

### Lentiviral Vector Generation

VSVG-pseudotyped lentiviral vectors encoding NGN2 were packaged as described previously (Naldini et al., 1996). Briefly, HEK-293T cells were plated onto 10-cm dishes the day before transfection at a density of 2.5x10^6^ cells/dish. 24 hrs later cells were co-transfected using calcium phosphate with 3.5 μg of envelope vector pMD2.G, 6.5 μg of packaging vector pCMVR8.74, and 10 μg of transfer vector pTet-O-Ngn2-puro. Transfections were done in Opti-MEM medium with complete exchange to mTeSR1 media after 24 hrs. Supernatants containing unconcentrated lentiviral particles were collected at 48 and 72 hrs, filtered through a 0.45 μm syringe filter, and stored at −80°C.

### Neural Differentiation and Characterization

For conversion to NSCs, iPSC clones on Geltrex were cultured in PSC Neural Induction Medium for 7 days, with media exchanges every other day. On day 7 cells were collected via Accutase and seeded onto fresh Geltrex-coated plates in Neural Expansion Medium (1:1 PSC Neural Induction Medium: DMEM:F12) with Y-27632 (5 μM). Medium was exchanged 24 hrs later to remove the ROCK inhibitor. Cells were propagated in culture for up to four passages, as NSC-like morphology typically improved over this period and any residual pluripotent-like cell colonies initially present would be depleted. To generate neurons, iPSCs were plated at clonal density on Geltrex in mTesR1 containing Y-27632 (10 μM). 24 hrs later, cultures were infected with NGN2-lentiviral vector by placing cells in undiluted inoculum + polybrene (8 μg/ml) for 4 hrs, followed by a medium exchange, and then a second round of infection the next day. 24 hrs after the second infection, cells were switched to DMEM:F12 medium + N2 supplement, NEAA, human brain-derived neurotrophic factor (BDNF; 10 ng/ml), human neurotrophin-3 (NT-3; 10 ng/ml) and doxycycline (2 μg/ml). The next day cells were treated with puromycin (1 μg/ml). After selection for 48 hrs, the neural population was collected via Accutase and seeded onto poly-D-lysin/laminin-coated plasticware in Neurobasal/Glutamax medium supplemented with B27, BDNF, NT-3, doxycycline, and Y-27632 (5 μM). The next day cells received a 50% medium exchange including additional selection with cytosine-β-D-arabinofuranoside (Ara-C) to deplete any non-neural cells resistant to puromycin. Ara-C was removed after 48 hrs, and cells continued to receive 50% media exchanges every other day until DIV 14.

NSC identity was confirmed by immunostaining for Nestin, Pax6, Musashi and Sox1, while neurons were labeled with antibodies against doublecortin, Tuj1, MAP2, TBR1, Tau, and VGLUT1. To further assess functional maturity of neurons, cells seeded during differentiation into 35-mm dishes (600K cells/dish) were loaded at DIV14 with the Ca^2+^ indicator dye, Fluo4-AM, at room temperature for 40 min, rinsed 3 times in PBS, then transferred to a Na^2+^-based extracellular solution containing (in mM): 140 NaCl, 5 KCl, 2 CaCl_2_, 1 MgCl_2_, 10 D-Glucose, 10 Hepes; pH 7.4. Cells were imaged using a Nikon Eclipse Ti microscope, Andor Zyla CMOS camera with a PE4000 Cool-LED light source. Exposure times were 40-60 ms and images were taken every 1 s. KCl (40 mM) or kainate (10 mM) were added for 10 s after 1 min baseline imaging recording. Individual cells were selected with Nikon software and resulting Ca^2+^ responses were calculated and graphed in Matlab as relative change in fluorescence intensity (ΔF/F).

### Strand-specific dUTP RNAseq Library Preparation

RNASeq libraries were prepared using TruSeq^®^ Stranded mRNA Library Kit (Illumina) and prepared per manufacturer’s instructions. In brief, RNA sample quality (based on RNA Integrity Number, RIN) and quantity was determined based on 2200 TapeStation and between 500-100 ng of total RNA was used to prepare libraries. 1 uL of diluted (1:100) External RNA Controls Consortium (ERCC) RNA Spike-In Mix (Thermo Fisher) was added to each sample alternating between mix 1 and mix 2 for each well in batch. PolyA bead capture was used to enrich for mRNA, followed by stranded reverse transcription and chemical shearing to make appropriate stranded cDNA inserts for library. Libraries are finished by adding both sample specific barcodes and adapters for Illumina sequencing followed by between 10-15 round of PCR amplification. Final concentration and size distribution of libraries were evaluated by 2200 TapeStation and/or qPCR, using Library Quantification Kit (KK4854, Kapa Biosystems), and multiplexed by pooling equimolar amounts of each library prior to sequencing. RNASeq libraries were sequenced to 40-100 million reads per library with 85-97% covered bases included in annotated mRNA.

### RNA Capture-Sequencing (CapSeq) Library Preparation

RNA CapSeq libraries were prepared using a combination of TruSeq^®^ Stranded mRNA Library Kit (Illumina) and SureSelectXT kit with a SureSelectXT custom capture library targeting a region of 400 kb on the X chromosome from *OGT* to *CXCR3*. In brief, cDNA was made from RNA using the TruSeq^®^ Stranded mRNA Library Kit (Illumina). 500-100 ng of total RNA was used to prepare libraries that were first PolyA-bead captured to enrich for mRNA, followed by stranded reverse transcription and chemical shearing to generate appropriate stranded ~175 bp length cDNA inserts. These cDNA inserts were end-repaired, adenylated, ligated to adapter oligos and then amplified with 5 cycles of PCR according to manufacturer’s instructions. After quantification, 750ng of each amplified DNA sample was hybridized overnight with the Capture Library. Following capture cleanup, each gDNA library was amplified with additional 16 cycles of PCR, which also tagged each sample with an index-specific barcode. Final products were quantified using the TapeStation 2200 and pooled for rapid mode sequencing on the Illumina. RNA CapSeq libraries were sequenced at the Broad Institute Genomic Services as 101bp paired-end reads on an Illumina HiSeq2000 platform. The experiment performed well, resulting in on an average 3401800 reads per sample, with average mapping rate of 86% and 15% of on-target rate of the sequenced reads.

### *De novo TAF1* Transcript Assembly and Quantification

For *de novo* assembly, alignments of each sample (both total RNAseq and RNA CapSeq if available) within the capture region were de-multiplexed, duplicate reads were removed, and reads pairs from the same cell types were then merged. Transcripts were assembled by Trinity v2.2.0 on each cell type (fibroblasts, NSCs, and neurons) with default settings (Grabherr et al., 2011). After the preliminary assembly, transcripts from the three cell types were compared and merged to a non-redundant list. These non-redundant *de novo* transcripts were then compared individually against human genome GRCh37.75 using BLAT to resolve internal splicing structures. For each transcript, we required the whole set of splice junctions to be supported by more than half of the RNAseq samples of any category combining cell types and genotype (controls, carriers and probands), or the *de novo* transcript was disregarded.

The abundance of the remaining transcripts was measured in each total RNAseq sample using RSEM v1.2.31 and bowtie2 v2.1.0 with the estimation of read start position distribution (“--estimate-rspd” option) (Li and Dewey, 2011). In parallel, gene counts of each total RNAseq sample were generated using HTSeq (version 0.6.1) by considering only fragments aligned to gene exons. The normalization factor across samples was estimated by DESeq2 (Love et al., 2014), by which we then calculated the overall normalized expression of *TAF1* in each sample. The absolute expression of each TAF1 isoform in a sample was calculated as the product of abundance of each transcript and overall normalized *TAF1* expression in that sample.

### Intron Retention Analysis

The approach according to Middleton et al. (2017) was applied to estimate genome-wide intron retention in each sample (Middleton et al., 2017; Wong et al., 2013). In general, the intronic expression levels were represented by the median depth of each intronic region without considering the pre-calculated low mappability regions and any intronic region that overlaps with expression features such as microRNAs and snoRNAs. The intron retention level of each intron was then estimated as a ratio of the intronic expression to the number of reads that connect the flanking exon junctions.

### CRISPR/Cas9 Nuclease-mediated Genome Editing

Cloning and characterization of sgRNAs targeting sequences 5’ and 3’ of the SVA were done as reported previously (Hendriks et al., 2015). Briefly, 20-nucleotide oligo sequences preceding *S.pyogenes* Cas9 (SpCas9) NGG PAM sites were designed immediately 5’ and 3’ of the SVA using Geneious DNA analysis software. *BbsI* overhang-containing oligos were synthesized for ligation into a *BbsI*-linearized pGuide sgRNA expression vector under control of the U6 promoter. After ligation, bacterial transformation and miniprep plasmid DNA isolation, sgRNA vectors targeting the SVA were verified by Sanger sequencing. Sequence-verified sgRNA plasmids (250 ng) and human codon-optimized pCas9-GFP (750 ng), containing SpCas9 nuclease fused to GFP under control of a CAG promoter, were transfected into HEK293T cells via Lipofectamine 3000 as recommended. Cells were maintained in culture for 72 hrs, and gDNA was isolated and subjected to PCR with primers amplifying a segment spanning the targeted region. PCR amplicons were purified and Sanger sequenced to confirm the presence of double-stranded DNA breaks (DSBs). Cleavage efficiency based on Sanger sequencing traces was estimated using TIDE (Tracking Indels by Decomposition) (Brinkman et al., 2014).

From that analysis, two guides targeting sites flanking the SVA insertion site were used to excise the retrotransposon from XDP iPSC clone, 33363-D. Cells grown to 70-80% confluence on Geltrex were collected via Accutase, triturated to single cells in Opti-MEM media, and transfected in suspension with 1.5 μg pCas9-GFP, 0.75 μg of each SVA sgRNA-encoding pGuide, 5 μL P3000 reagent, and 3.75 μL Lipofectamine 3000. After 15 minutes, fresh mTeSR1 with Y-27632 (10 μM) was added to each reaction, and cells were plated on Geltrex. A complete medium exchange was performed the next day to remove ROCK inhibitor, and cells were maintained for an additional 48 hrs. GFP-positive cells were then collected as single cells via Accutase in 250 - 750 μL DPBS with Y-27632 (10 μM), filtered through a 35-μm mesh cell strainer, and sorted on a BD FACSAria™ Fusion SORP Cell Sorter using a 100 μm nozzle at a pressure of 20 psi. Sorted cells were collected and plated at clonal density of 2.5 - 3.0 x 10^4^ cells per 10-cm dish in mTeSR1 with Y-27632 (10 μM) and MycoZap™ (1:250 dilution). A complete medium exchange was performed the next day to remove ROCK inhibitor and every day thereafter until single cell-derived colonies reached 2-4 mm in size. Single colonies were manually picked into individual wells of a 96-well plate, propagated until approximately 90% confluent, then collected using Accutase and divided into two 96-well companion plates: one to which freezing media (80% FBS/20% DMSO) was added 1:1 to each well and stored at −80°C (clone recovery plate); the other kept in culture with cells propagated until reaching approximately 90% confluency again. Cells were then lysed overnight in 50 μL lysis buffer (10 mM Tris-HCl, pH 7.5-8.0; 10 mM disodium EDTA; 10 mM NaCl; 0.5% [w/v] sarcosyl) at 56°C. The next day gDNA was precipitated with ice cold ethanol (95% v/v) for 2 hr at −20°C, pelleted, washed with ethanol (70% v/v), and resuspended in 30 μL ddH2O + 0.1 mg/mL RNase A. Samples were screened by PCR with primers amplifying a segment spanning the SVA insertion site, and successful excision of the SVA was detected based on the size of the amplicon (~0.6 kb vs. 3.0 kb with or without the SVA, respectively) determined by electrophoresis. Positive amplicons lacking the SVA were subsequently confirmed by Sanger sequencing. Successfully edited clones were then recovered from the sister plate and propagated as described above, confirmed to retain normal karyotype and expression of pluripotency markers, and differentiated to NSCs and NGN2-neurons.

### Data Processing and Computational Analyses for RNA CapSeq and Total RNAseq

Read pairs of RNA CapSeq and RNAseq were trimmed by trimmomatic v0.36 using Illumina Truseq adapters and primers. The trimmed read pairs were then aligned to human genome (GRCh37, Ensembl release 75) with SVA insert at position X:70660363 by STAR 2.4.2a (Dobin et al., 2013), allowing 5% mismatches with a unique mapping.

To identify factors influencing gene counts in each cell type, PCA analyses were performed on log-transformed pseudocounts (log2[count+1]) normalized by sample-specific sequencing depth. Hierarchical clustering of the samples based on Euclidian distance of all principal components with complete linkage and subsequent dynamic branch cutting was used for identification of PCA clusters and was implemented using R packages fastcluster (Mullner, 2013) and dynamicTreeCut (Langfelder et al., 2008) correspondingly. To estimate log2-fold changes in XDP vs. control samples, generalized linear models were used to model counts by negative binomial distribution using R package DESeq2 (v 1.10.1) (Love et al., 2014), where PCA cluster and Genotype were included as factors in the model design: counts~PCA+Genotype. Estimated log2-fold changes were tested for significance using Wald test, and corresponding *p*-values were adjusted for multiple hypothesis testing using Benjamini-Hochberg adjustment method (pBH). Genes were called significant if corresponding pBH<0.1.

### Co-expression Networks, Gene Ontologies and Pathways Analysis

Enrichment for gene ontologies was tested using R package topGO (version 2.22.0), using only curated gene ontology assignments (excluding evidence with codes ND, IEA, NR) with algorithm “weight01.” Co-expression network analysis was performed using R package WGCNA (Langfelder and Horvath, 2008) for each cell type separately using unsigned network type. Only genes with median absolute deviation (MAD) >0 and with counts > 10 in at least half of the samples within a cell type were considered. Variance stabilizing transformation was applied to counts prior to co-expression module detection. Soft power was selected such that the scale-free topology fit (R^2^) > 0.8. Merge of the modules with similar eigengenes was performed if the number of modules was > 30. Module membership for each gene was re-evaluated based on the module membership *p*-value; if *p*-value > 0.01, the gene was marked as unassigned (Module 0). Correlation of modules to transcript expression was calculated using Pearson’s correlation. Correlation of modules to intron retention was calculated using Spearman’s correlation.

